# Partially-functional exhausted CD8+ T cells can contribute to short-term viral suppression: a computational prediction for children with perinatal HIV

**DOI:** 10.1101/2025.09.18.677001

**Authors:** Alexis Hoerter, Alexa Petrucciani, Fatma Marshed, Mussa Mwamzuka, Aabid Ahmed, Alka Khaitan, Elsje Pienaar

## Abstract

We and others have reported evidence of T cell exhaustion in children with perinatal HIV with increased expression of inhibitory receptors PD-1, CD160, and TIM-3, but there is limited data on the virologic functional consequences of this immune exhaustion. We address this by using an immune database from Kenyan children with perinatal HIV and unexposed controls. We computationally integrate T cell profiles of differentiation, activation and exhaustion in an agent-based model (ABM) to predict how T cell exhaustion impacts viral control following HIV exposure in vitro. Our ABM includes macrophages, CD4 and CD8 T cells, cytokines, and HIV. Model mechanisms include viral dynamics, macrophage activation, T cell activation and proliferation, cytotoxic T cell killing, and cytokine/HIV diffusion and degradation. Participants are grouped by HIV plasma viremia and by age, less than 5 years or 5-18 years. Our findings indicate that cells from virally active participants, who have the highest levels of exhaustion, have lower predicted viral concentrations and infected cells compared to other participant groups during new infection. However, this coincides with higher cell death, suggesting that short-term viral control is associated with excessive inflammation, which could be detrimental long-term. Cells from virally suppressed participants older than 5 years can maintain lower viral concentrations while limiting cell death, reflecting a more sustainable short-term immune response. In virally suppressed children younger than 5 years, immune response patterns strongly resemble the age-matched healthy control group, suggesting early viral suppression may preserve antiviral immune responses. Our model predicts unique patterns of cell death for each participant group, with CD8 T cell death being dominant in virally active groups and CD4 T cell and macrophage death being dominant in healthy and virally suppressed groups. Finally, exhausted CD8 T cells are predicted to contribute significantly to CD8 T cell killing, proliferation, and activation in the virally active group, indicating partially functional CD8 T cells can still contribute to short-term viral control. Our analysis functionally integrates participant-specific immunophenotypic data to allow quantification of the extent, mechanisms, and impact of immune dysfunction in perinatal HIV and could inform pediatric HIV remission and cure strategies.

**Author Summary:** Cytotoxic CD8+ T cells are vital for our ability to kill infected cells. However, chronic stimulation of CD8+ T cells can result in T cell exhaustion, leading to cellular dysfunction and reduced ability to fight infections. Here we ask: What impact does CD8+ T cell exhaustion have on the ability to control HIV replication over the short term?

We answer this using a computational model, anchored in experimental data from children with perinatal HIV. We use participant-specific data on dozens of immune markers that characterize the state of their immune system. We carefully map these markers to known immune cell functions and use our model to predict how immune cells with these participant-specific markers would respond to HIV infection.

Our results suggest that earlier viral suppression can lead to better immune function in children with perinatal HIV. Our findings also suggest that exhausted CD8+ T cells could still contribute to fighting HIV infection, but only if there are enough of them to compensate for their reduced functionality.

Our work uses computational models to put valuable participant-specific data into context, allowing us to predict short-term infection outcomes and better understand immune function in children with perinatal HIV.

## 1. Introduction

Despite effective antiretroviral therapy (ART) of both mothers and children, 160,000 children are infected with HIV each year(1). As in adults, ART can effectively control, but not eliminate, HIV in children. Long-term infection in children is also associated with chronic inflammation leading to T cell exhaustion and weakened immune responses to HIV. However, the extent, mechanisms and impact of this immune dysfunction in children with perinatal HIV (CPHIV) remains unclear. Immune exhaustion, and CD8 T cell dysfunction in particular, has been identified in CPHIV through elevated expression of immune inhibitory receptors (IRs) such as PD-1, TIM-3, 2B4 and CD160. We and others have reported elevated PD-1 levels on CD8 T cells in CPHIV, but there is limited data on other IRs and their functional consequences(2–5). In a South African child with sustained virologic remission, the one distinct immune feature was high PD-1 levels, warranting further investigation into PD-1 as a biomarker of clinical outcomes or treatment interruption in children(6).

Collection and analysis of pediatric samples to answer these questions are challenging. Pediatric study participants are more difficult to recruit than adult participants(7). If samples can be obtained from children, the smaller amounts of blood that can be safely and feasibly collected from children limit the types of analyses that are possible. For example, functional assays such proliferation assays can only be performed on total PBMCs since the small sample volumes are insufficient to sort immune subsets. These bulk analyses therefore also pose an analysis challenge in quantifying the relative contribution of individual immune cell subtypes.

Here we address these cohort building and data analysis challenges by pairing clinical parameters, immune phenotypic data and *in vitro* functional assays with computational systems biology approaches. We leverage an established database of clinical and immunologic parameters from a cohort of CPHIV and a control group of children unexposed to HIV (CHU) (3,5,8–13). We couple these participant-specific data with a mechanistic computational model that emulates *in vitro* HIV infection assays. Our established agent-based models (ABMs) (14–16) provide virtual versions of *in vitro* HIV infection assays, incorporating viral, innate and adaptive immune dynamics. These computational models, parameterized with participant-specific data, complement their experimental counterparts by allowing quantitative characterization of complex host-pathogen interactions with high spatial and temporal resolution.

Here, we use our computational models to predict the impact of exhausted CD4 and CD8 T cells (expressing PD-1, TIM-3, and/or CD160) on short-term viral control and immune responses to new HIV exposure. Our simulations provide testable predictions and generate new hypotheses about the functional impact of T cell exhaustion on HIV-specific immune responses. Our findings serve to integrate complex clinical and immune datasets, predict host responses to viral blips and inform future HIV functional cure approaches.

## 2. Materials and Methods

### 2.1 Cohort and Immune Phenotypic Data

We utilize an established database from the Pediatric Immune Activation (PIA) study as described previously(3,5,8–13). Briefly, a total of 242 children between the ages of 2 months to 20 years were enrolled from Bomu Hospital, an HIV-centered nonprofit hospital in Mombasa, Kenya from 2011-2012. The cohort was comprised of 156 CPHIV, including 76 children who were treatment naïve, and 84 children who had been on antiretroviral therapy for at least six months. As controls, 82 children unexposed to HIV (CHU) were enrolled.

Blood samples were collected from all children to perform HIV viral loads and CD4 cell counts and to isolate and cryopreserve PBMCs and plasma. For each participant, we performed a broad immune phenotype analysis by flow cytometry to delineate innate (monocytes, dendritic cells, natural killer cells) and adaptive immune subsets (B, Th1, Th2, Th17, Th22, regulatory T cells), T cell differentiation states (naïve, central, transitional and effector memory and EMRA), chemokine receptor expression, T cell activation (CD38, HLA-DR) and exhaustion (PD-1, CD160, TIM3, 2B4), and proliferative (Ki67) and cytotoxic (granzyme B, perforin) potential. Flow gating strategies are detailed in (3,5,8). The immune phenotypic data was compiled with clinical data including complete blood counts, HIV viral load, CD4 T cell counts, and antiretroviral treatment to build a master de-identified database of clinical and immune parameters. In total, 116 participants have complete immune phenotypic profiles for simulation parameters discussed below in Section 2.2.9 and are used in simulations. CPHIV was divided into groups based on viral suppression for analysis in our computational models of HIV infection dynamics. CPHIV virally suppressed (CPHIV-VS) participants are defined as those with HIV viral load less than 200 copies/mL and CPHIV virally active (CPHIV-VA) are defined as those with viral loads equal to or above 200 copies/mL. **Table 1** shows the demographic and clinical characteristics of the participants used in simulations.

**Table 1:** Demographic and clinical characteristics of participants with immune profiles used in simulations.

|  | Age 0-5 years |  |  |  | Age 5-20 years |  |  |  |
| --- | --- | --- | --- | --- | --- | --- | --- | --- |
|  | CHU | CPHIV-VS | CPHIV-VA | p-value | CHU | CPHIV-VS | CPHIV-VA | p-value |
| n | 12 | 13 | 24 |  | 20 | 19 | 28 |  |
| Age (years)<sup>a</sup> | 2.6 (1.5 – 3.3) | 3.2 (1.7 – 3.6) | 2.4 (1.5 – 3.0) | NS<sup>b</sup> | 13.2 (10.6 – 17.3) | 11.84(8.3 – 13) | 11.7 (9.1 – 16.5) | NS<sup>b</sup> |
| Female sex, No. (%) | 10 (83.3%) | 8 (61.5%) | 9 (37.5%) | p<0.002<sup>c</sup> | 4 (20%) | 10 (52.6%) | 16 (57.1%) | p=0.01<sup>c</sup> |
| HIV copies/mL<sup>a</sup> |  | 110 (110-110) | 65725 (3099 – 257400) | p<0.001<sup>b</sup> |  | 110 (110-110) | 52058 (6738 – 172700) | p<0.001<sup>b</sup> |
| Absolute CD4 count (cells/mm<sup>3</sup>)<sup>a</sup> | 1595 (1314 – 2116) | 1703 (1220 – 2194) | 1552 (1065 – 2021) | NS<sup>b</sup> | 779 (591 – 931) | 957 (674–1216) | 498 (313 – 790) | p<0.001<sup>b</sup> |

(a) Median values with interquartile range
(b) Kruskal-Wallis test
(c) Chi-squared test
(NS) not significant; Significance determined at the threshold of p-value < 0.05

### 2.2 Simulation Description

Our ABM uses unique, individual participant immune phenotypic data from the cohort described in Section 2.1 to predict the immune response to new HIV infection *in vitro*. For simulations, we group participant data by HIV RNA copies/mL at the time of data collection: HIV RNA copies/mL = 0 (CHU), < 200 (CPHIV-VS), and ≥ 200 (CPHIV-VA) as well as by age (0-5 years, and ≥ 5 years). We use the same model rules as described in (14–16), and briefly summarized below, but with the initial parameters described below as well as mechanism changes to reflect HIV infection. The ABM consists of macrophages, CD4 and CD8 T cells, two diffusible cytokines TNFα and IFNγ, and HIV. All cell types have their own rules that are executed simultaneously mimicking realistic cell behavior. General model mechanisms are described in **Fig. 1**. A detailed flowchart of possible clinical markers for T cells is given in **Fig. 2**. An outline of the general clinical markers mapped to simulation mechanisms is given in **Table 2**, and model parameters are given in **Table 3**. We simulate an *in vitro* HIV infection for 14 days with a timestep of 6 minutes, approximately the time it takes for a macrophage to move its own length (17–20) (20 μm) or one grid square(14). The simulation is implemented in Java using Repast Simphony (21). MATLAB and Python are used for data analysis and visualization. We use ChatGPT to create and edit graphing scripts for MATLAB, and to summarize methods text from (14–16) for the model mechanisms that are not changed in this work. Any content generated by ChatGPT was carefully reviewed by our team for accuracy.

**Fig. 1:**
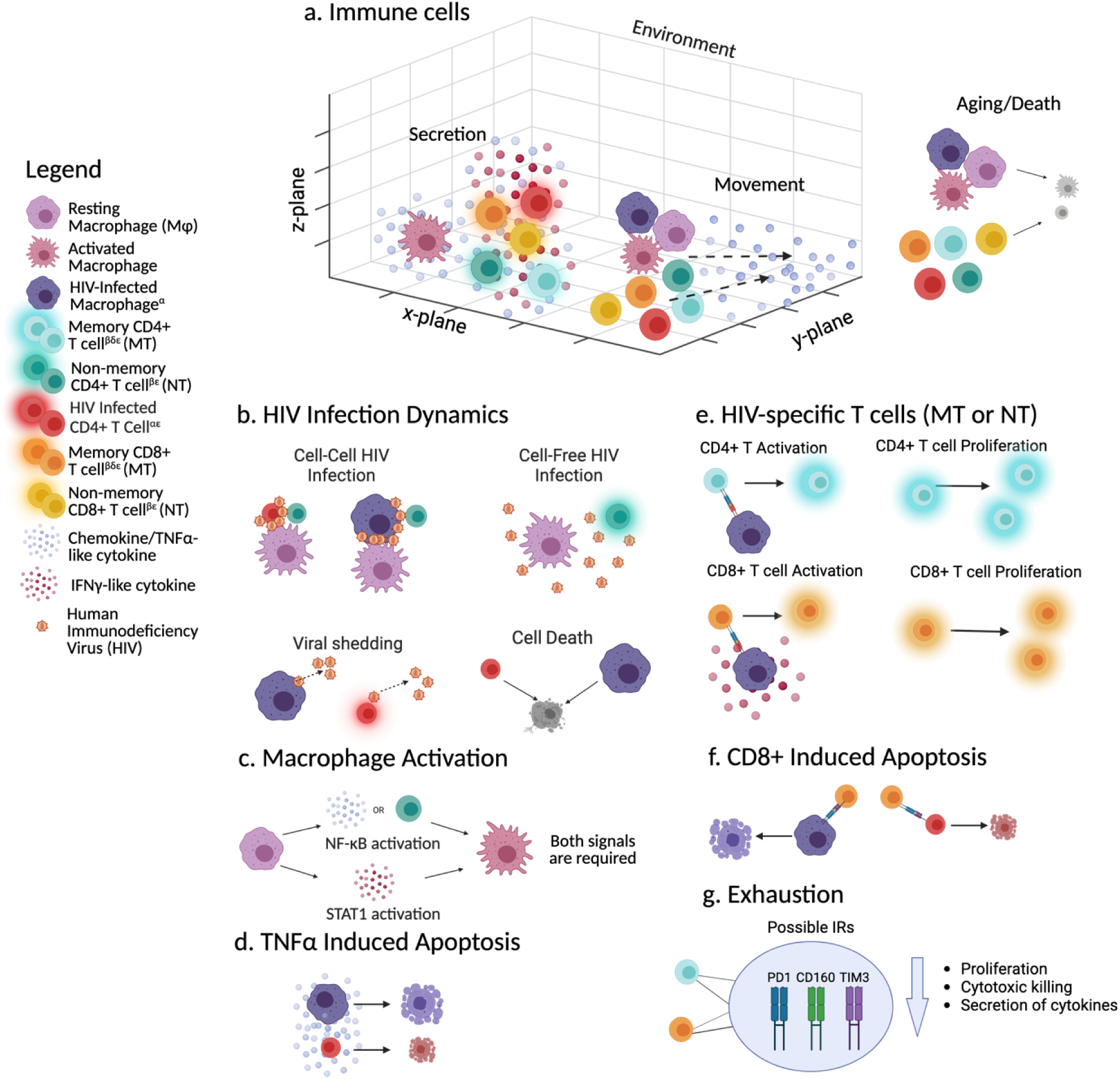
Schematic of general ABM agents and rules adapted from(14–16). CD4 and CD8 T cells are divided into two main populations: memory and non-memory. a) All immune cells exist in a 3D environment and remain at the bottom z-layer unless on top of another cell. Each compartment can hold only one immune cell. Only activated or recently differentiated CD4 T cells and CD8 T cells secrete TNFα and IFNγ (cell must be TNFα/IFNγ+) and activated macrophages secrete TNFα. All immune cells move probabilistically along the TNFα concentration gradient. All immune cells have ages (resting and activated) and die when their maximum lifespan is reached. b) HIV can infect activated and CCR5+ CD4 T cells or macrophages either via cell-free concentration (HIV >1) or through infected cell contact. Only activated HIV-infected cells shed virus. After a maximum infection time, HIV-infected cells die. c) Macrophages can become activated by activating both NF-κB and STAT1 pathways. d) Any cells can apoptose based on the local concentration of TNFα. e) HIV-specific CD4 and CD8 T cells can become activated/differentiated and can divide. CD8 T cells need the HIV-infected cell to be STAT1 activated. f) CD8 T cells can kill (apoptose) HIV-infected cells (if memory must be HIV-specific and Granzyme+ or Perforin+; if recently differentiated effector must be HIV-specific. g) Up to three IRs (PD1, CD160 or TIM3) can be found on memory CD4 (blue) and CD8 T (orange) cells that can decrease their proliferation probabilities, cytotoxic killing and secretion of cytokines (IFNγ and TNFα). The severity of the loss of function is determined by the number of IRs on the cell. α: Must be CCR5+; β: Can be HIV-specific; δ: Can have IRs; ε: Glow represents activated cell (memory) or recently differentiated effector cell (non-memory); +: Only activated cells at the start can perform any functional operations. After initialization, only HIV-specific cells can perform functions once activated. See Figure 2 for flowchart of possible markers that can be found on T cells. Created in Biorender. Created in BioRender. (2025) https://BioRender.com/t09v629

**Fig. 2:**
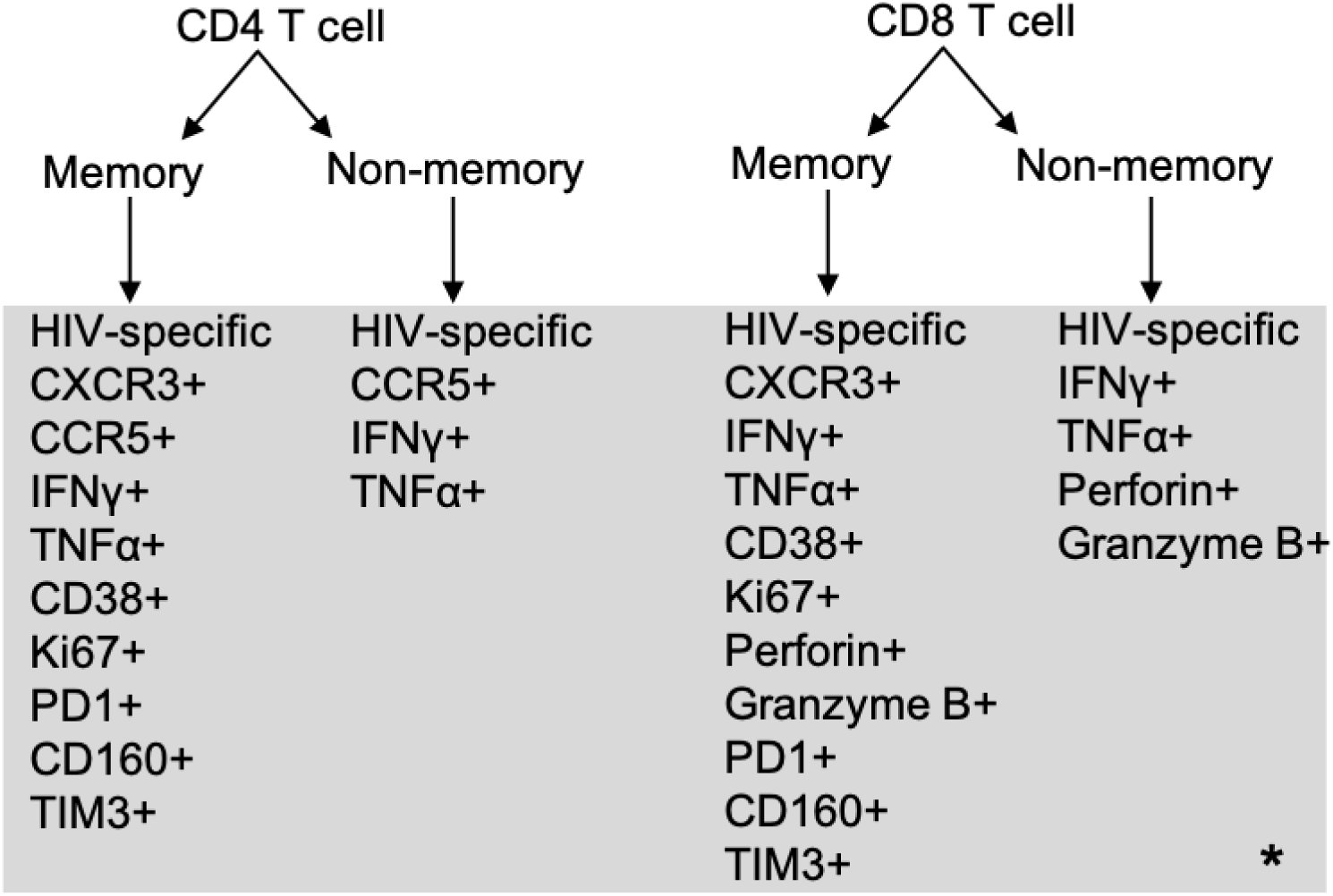
Flowchart of the various clinical markers that can be assigned to CD4 and CD8 T cells in our simulation. *: the markers in the grey box are randomly assigned in the simulation to their given population, resulting in a simulated population of cells with a heterogeneous total number of markers per cell. Meaning any given cell can have any number of the markers listed under its respective subtype. It’s important to note that if a memory non HIV-specific T cells starts activated (Ki67+ or CD38+) then it can perform cytokine secretion (if IFNγ+ and/or TNFα+), help activate macrophages and kill infected cells if a CD8 T cell (only if granzyme B+ or perforin+) otherwise it does nothing but move regardless if it is positive for other markers. Non-memory HIV-specific cells can become activated by interacting with HIV infected cells, but none start activated. These cells can also be positive for other markers, but if they are not HIV-specific they will never be activated to perform any of the functions those markers are associated with.

**Table 2:** Mapping of clinical markers to simulation mechanisms.

| Clinical Marker | Simulation mechanism |
| --- | --- |
| CXCR3+ | T cell has a greater individual probability of moving |
| CCR5+ | CD4 T cell can be infected by HIV |
| HIV-specific | Antigen specific T cell that can perform actions based on other markers (activate macrophages, proliferate) |
| IFNγ+<br>TNFα+ | T cell can secrete these cytokines |
| CD38+ | T cell will be activated at start of simulation |
| Ki67+ | T cell will be activated at start of simulation and has the ability to proliferate |
| Perforin+<br>Granzyme B+ | CD8 T cell that has the ability to kill HIV-infected cells |
| PD1+<br>CD160+<br>TIM3+ | Inhibitory markers (IRs)/exhaustion markers on T cell |

**Table 3:** Original parameter ranges used to identify 20 representative parameter sets for a robust HIV infection. Parameter ranges were determined based on available literature or defined based on biological feasibility if no literature values could be found and then estimated using LHS.

| Name | Value | Units | Reference |
| --- | --- | --- | --- |
| **CD4 Parameters affected by IRs** |  |  |  |
| ActivatedCD4IFNSecretion* | 5-40 | Molecules/cells/s | (40,41) |
| ActivatedCD4TNFSecretion* | 5-40 | Molecules/cells/s | (40,41) |
| proliferationProbabilityCD4* | 0.1-1 | Per timestep |  |
| **CD8 Parameters affected by IRs** |  |  |  |
| ActivatedCD8IFNSecretion* | 5-40 | Molecules/cells/s | (40,41) |
| ActivatedCD8TNFSecretion* | 5-40 | Molecules/cells/s | (40,41) |
| CD8HIVSpecificKillProbability* | 0.1-1 | Per timestep |  |
| proliferationProbabilityCD8* | 0.1-1 | Per timestep |  |
| **HIV Parameters** |  |  |  |
| startingFreeVirus* | 1-100,000 | Concentration/simulation volume | e |
| eclipsePhaseForVirusProduction | 24 | hours | (42) |
| eclipsePhaseForVirusProductionVariance | 0.2 |  |  |
| hivCellCellInfectionProbability* | 0.25-1 | Per interaction | e |
| hivFreeVirusInfectionProbability* | 0.25-1 | Per interaction | e |
| hivProductionRateVariance | 0.1 |  |  |
| maxHIVInfectedCellLifespan | 2.2 | days | (38) |
| maxHIVInfectedCellLifespanVariance | 0.2 |  |  |
| viralProductionRatePerDayCD4 | 5 | $10^4$ virions/day | (43) |
| viralProductionRatePerDayMacro | 5 | $10^4$ virions /day | (43) |
| virusDegradationRate | 2.67 | $10^{-5} s^{-1}$ | (38) (free virions in plasma lifespan of 0.3 days turned into rate) |
| virusDiffusionCoefficient | 4.38 | $10^{-8} cm^2/s$ | (44) (Stokes Einstein #) |
| **CD4 Parameters not affected by IRs** |  |  |  |
| activatedMovementProbabilityCD4*+ | 0.1-1 | Per timestep | e |
| baseMovementProbabilityCD4*+ | 0.1-1 | Per timestep | e |
| CD4ActivationProbability* | 0.1-1 | Per interaction | e |
| CD4DeactivationProbability* | 0-0.2 | Per timestep |  |
| cd4PopulationDoublingTime* | 4-16 | Hours | (45–47) |
| cd4PopulationActivatedLifespan | 4 | Days | (45–48) |
| cd4PopulationMaxLifespan | 340 | Days | (49) |
| cd4DoublingTimeVariance | 0.1 |  |  |
| cd4LifeSpanVariance | 0.1 |  |  |
| maximumCD4Generations* | 4-10 | Generations | (45–47,50,51) |
| hivSpecificNaiveCD4Fraction | 0.02 |  |  |
| hivSpecificMemoryCD4Fraction | 0.02 |  |  |
| percentCCR5NaiveCD4s* | 0.021-0.83 |  | (52) |
| **CD8 Parameters not affected by IRs** |  |  |  |
| activatedMovementProbabilityCD8*+ | 0.1-1 | Per timestep | e |
| baseMovementProbabilityCD8*+ | 0.1-1 | Per timestep | e |
| CD8HIVSpecificActivationProbability* | 0.2-1 | Per interaction | e |
| CD8HIVSpecificDeactivationProbability* | 0-0.2 | Per timestep |  |
| cd8PopulationDoublingTime* | 4-16 | Hours | (45–47) |
| cd8PopulationMaxActivatedLifespan | 4 | Days | (45–48) |
| cd8PopulationMaxLifespan | 340 | Days | (49) |
| cd8DoublingTimeVariance | 0.1 |  |  |
| cd8LifeSpanVariance | 0.1 |  |  |
| maximumCD8Generations* | 4-20 | Generations | (45–47,50,51) |
| hivSpecificCD8Fraction | 0.02 |  |  |
| hivSpecificCD8NaiveFraction | 0.02 |  |  |
| **Macrophage Parameters** |  |  |  |
| ActivatedMacrophageTNFSecretion* | 5-40 | Molecules/cell/s | (40,41) |
| activatedMovementProbabilityMacro* | 0.1-1 | Per timestep | e |
| baseMovementProbabilityMacro* | 0.1-1 | Per timestep | e |
| macrophageLifeSpanVariance | 0.1 |  |  |
| macrophagePopulationMaxActivatedLifespan | 10 | Days | (28) |
| macrophagePopulationMaxLifespan | 100 | Days |  |
| nfkBspan* | 0.11-115 | Hours | (29) |
| nfkBVariance | 0.1 |  |  |
| stat1Span* | 0.11-115 | Hours | (29) |
| stat1Variance | 0.1 |  |  |
| TNFthresholdForNFkBActivation* | 10-500 | Molecules | e |
| IFNthresholdForStat1Activation* | 10-500 | Molecules | e |
| percentCCR5Macs* | 0.242-0.902 |  | (52) |
| activatedMacrophageProportion* | 0.05-0.5 |  |  |
| **Cytokine/General Immune Cell Parameters** |  |  |  |
| InitialPBMCs* | 6000-10000 |  | e |
| IFNDegradationRatePerSecond* | 0.96-10 | s<sup>-1</sup> | (53–57) |
| IFNDiffusionCoefficient* | 0.1-1 | cm<sup>2</sup>/s | (53–55,57–59) |
| TNFDegradationRatePerSecond* | 0.96-10 | s<sup>-1</sup> | (53–57) |
| TNFDiffusionCoefficient* | 0.1-1 | cm<sup>2</sup>/s | (53–55,57–59) |
| TNFthresholdForImmuneCellMovement* | 10-500 | Molecules | e |
| Kapop* | 9.93E-05 -<br>0.0099336<br>5 |  | e |
| Kd* | 0.5450045<br>-<br>54.500450<br>3 | 10<sup>6</sup> | e |
| tauapop* | 0.06 -<br>0.24 |  | e |

\*: 20 parameter sets determined from the initial LHS range. +:parameter is affected by CXCR3; e: no literature value could be identified therefore this parameter was estimated to be biologically feasible and estimated through LHS.

#### 2.2.1 Environment

The ABM environment is a 3D grid with 100 x 100x 10 compartments, representing a 2 mm x 2 mm x 0.2 mm subsection of a well-mixed *in vitro* HIV infection assay (e.g. a well in a 96 well plate). Each grid compartment is a cube with a width of 20 μm, with toroidal (x-y plane) and no flux (z plane) boundaries (**Fig. 1a**).

#### 2.2.2 Cytokines

We represent IFNγ and TNFα as continuous variables and we numerically solve partial differential equations describing their diffusion and degradation using a 3D alternating-direction explicit method (14,22,23).

#### 2.2.3 Immune Cells

All immune cells (macrophages, CD4 and CD8 T cells) move by following a TNFα chemotactic gradient, and can “fall” due to simulated gravity. Movement is probabilistic and based on cells’ current functional state, resting or activated (14–16). In brief, cells will generally move towards higher concentrations of TNFα. The concentration of TNFα in each grid in the surrounding Moore neighborhood are summed up and then each grid’s individual TNFα concentration is divided by the sum of the concentration in the Moore neighborhood to give a probability of moving to that grid square. See (14) for more details regarding movement rules. Cell aging accelerates upon activation, and TNFα-induced apoptosis is simplified from (24) and described in Equation 1 in Supplement Materials.

#### 2.2.4 Macrophages

HIV-infected macrophages, representing all antigen-presenting cells, activate HIV-specific T cells, approximating MHC II and I antigen presentations(25). Macrophages secrete TNFα upon activation through both NF-κB, induced by TNFα or CD4 T cell contact, and STAT1 pathways induced by IFNγ (26–29).

#### 2.2.5 T cells

CD4 and CD8 T cells in the simulation are divided into 2 subtypes: memory and non-memory. We describe the characteristics of each subtype below, along with the markers from the immune phenotypic database that are used to parameterize these subtypes in the model. **Table 2** provides a mapping of clinical markers to simulation mechanisms and **Fig. 2** shows a flowchart of the possible clinical markers that can be found on simulated T cells. These clinical markers were determined for certain cell populations and not for their single-cell combination, therefore the combinations of markers per cell is random. **Table 1** in Supplement Materials provides a more detailed clinical marker to simulation parameter mapping.

##### 2.2.5.1 Functions assigned to memory CD4 T cell phenotypes

Memory CD4 T cells in the simulation are characterized by expression of immune markers for movement (CXCR3+), infection susceptibility (CCR5+), cytokine secretion (IFNγ+, TNFα+), activation (CD38+), proliferation (Ki67+), HIV specificity, and inhibitory receptors (PD1+, CD160+, and TIM3+ (Section 2.2.7)) based on the immune phenotypic data. The chemokine receptor CXCR3 has been identified to have an impact on cellular movement and CXCR3+ cells are 5-10 fold more efficient at moving (30,31). To keep things as simple as possible, all cells are initially assigned base and activated movement probabilities according to our estimated parameter ranges. If a cell is CXCR3+ then a random number between 5 and 10 is sampled and multiplied to the base and activated movement probabilities. Equation 2 in Supplement Materials demonstrates how this is calculated for a cell. This results in most CXCR3+ cells being guaranteed to move at every timestep. Proliferated cells that are CXCR3+ will sample new movement probabilities when they are created. CCR5+ cells represent cells that are susceptible to HIV infection as this is one of the main coreceptors necessary for HIV entry into cells. Only cells positive for IFNγ will secrete IFNγ if activated. Likewise, only cells positive for TNFα will secrete TNFα if activated. We use the CD38+ marker to determine the number of cells that start the simulation already activated. We use the Ki67+ marker to determine the number of cells that are already at some random point in their proliferation cycle and also activated. For the purpose of our simulation, we assume that CD38+ and Ki67+ cells represent different pathways of activation and thus do not overlap these markers. Therefore, only the memory cells that are Ki67- will be allowed to be CD38+. Cells that start with the Ki67 marker will have their current “generation” sampled between (1,maximumCD4Generations). HIV-specific CD4 T cells can become activated by randomly selecting and interacting with a macrophage near them. If the macrophage is HIV-infected, the T cell has some probability of becoming activated, reflecting MHC II antigen presentation(25). Activation is not for an indefinite time period; CD4 T cells have a chance to deactivate each timestep which is an approximation for anti-inflammatory mechanisms. Any activated CD4 T cell can replicate within a variable timeframe up to a generational limit (maximumCD4Generations) and will secrete IFNγ and TNFα at defined rates if positive for those markers. Activated CD4 T cells can also help macrophages get NF-κB activated [see [(14–16) for more details]. Memory CD4 T cells can have 0, 1, 2, or 3 IRs, which impact their other functional abilities (discussed below in 2.2.7). The frequency of these markers is determined by individual participant data, but co-expression of all these markers is randomly assigned per cell during simulation initialization due to a lack of combination staining. Resting HIV-specific and non-specific CD4 memory T cells move around the simulation without further action unless they are CCR5+, in which case they can be infected. The only action non-specific CD4 memory T cells can perform is cytokine secretion but only if activated from the beginning.

##### 2.2.5.2 Functions assigned to memory CD8 T cell phenotypes

Memory CD8 T cells are characterized by expression of immune markers for movement (CXCR3+), cytokine secretion (IFNγ+, TNFα+), activation (CD38+), proliferation (Ki67+), cytotoxic killing (perforin+, granzyme B+), HIV specificity, and exhaustion (PD1+, CD160+, and TIM3+ (Section 2.2.7)) based on the immune phenotypic data. We assume the percentage of CXCR3+ memory CD8 cells is similar to that of CD4 memory cells, due to lack of data for the CD8 T cell population. The framework for CXCR3+, IFNγ+, TNFα+, CD38+, and Ki67+ for memory CD4 T cells (see 2.2.5.1) applies to memory CD8 T cells as well. Memory CD8 T cells that are either perforin+ or granzyme B+ act as cytotoxic T cells, and if they are HIV-specific, can probabilistically kill HIV-infected cells. HIV-specific CD8 T cells can become activated by finding an HIV-infected macrophage that is also STAT1 activated which acts as a proxy for co-stimulation(32) reflecting MHC I antigen presentation(25). Activated HIV-specific CD8 T cells can kill HIV-infected cells (CD4 T cell or macrophage) with some probability, secrete TNFα and IFNγ if positive for those markers, and proliferate in the same way as CD4 T cells. Memory CD8 T cells can have 0, 1, 2, or 3 IRs which impact their other functional abilities, if also positive for those markers, which is discussed below in 2.2.7. The frequency of these markers is determined by individual immune phenotype, but the co-expression of all these markers is randomly assigned per cell during simulation initialization due to a lack of combination staining. Resting HIV-specific and non-specific CD8 memory T cells move around the simulation without further action. The only action non-specific CD8 memory T cells can perform is cytokine secretion but only if activated from the beginning.

##### 2.2.5.3 Non-memory T cell

T cells that are not memory subtypes are considered naïve T cells at the start of the simulation with some portion being HIV-specific. Naïve CD4 and CD8 T cells with HIV-specific markers can be primed within the simulation upon encountering HIV-infected macrophages, turning into recently differentiated effector T cells. Additional immune markers in this recently differentiated effector T cell population include cytokine secretion (IFNγ+, TNFα+) and cytotoxic killing (perforin+, granzyme B+) (CD8 only). These recently differentiated effector T cells can secrete cytokines (only if positive for IFNγ or TNFα), proliferate, activate macrophages (CD4 only) and kill HIV-infected cells (only perforin+ or granzyme+ CD8s). These cells, which cannot become exhausted in the simulation, exhibit movement probabilities akin to CXCR3- cells. Their TNFα and IFNγ secretion rates, along with proliferation and killing probabilities (CD8), are determined as for non-exhausted cells (**Equation 1**). All other naïve T cells, move around the simulation without further action unless they are CCR5+ (CD4), in which case they can be infected.

#### 2.2.6 HIV Specificity

In a subset of participants, the frequency of HIV-specific T cells was identified by *ex vivo* proliferation assays as previously described(3,5). Since proliferation assays were not performed for all participants, we set the value of HIV-specific CD4 and CD8 T cells to be 2% for every participant.

#### 2.2.7 Markers of T cell Exhaustion

The immune database includes the frequency of PD1+, CD160+, and TIM3+ CD4 and CD8 T cells, however, the explicit function of each of these IRs is unclear. Prior studies show that CD8 T cell dysfunction or exhaustion increases with co-expression of multiple IRs(5,33). We assumed that the number of IRs displayed on each cell correlates linearly with the degree of exhaustion, at least for CD8 T cells (5). It is unknown whether this is true for CD4 T cells, but we will assume for our purposes here that it is similar. Therefore, we will employ a binning approach linking IR co-expression count to exhaustion severity, affecting functional parameters like proliferation rate and cytokine secretion in CD4/CD8 T cells, and CD8 T cell killing probability (**Fig. 1g**).**Equation 1** demonstrates how the number of IRs impacts the decrease in proliferation parameters through the use of a random multiplier. Secretion and killing parameters are determined in the same way. We assume IRs are inherited upon cell proliferation(33,34). A new multiplier for each parameter is randomly selected based on the number of IRs for the newly proliferated cell. Cells cannot become newly exhausted in our simulations as the time scale we are evaluating is too short.

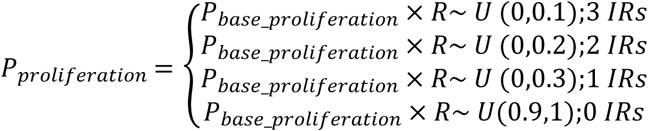

**Equation 1:** Assuming a linear correlation with the number of IRs expressed on a cell with more severe exhaustion, we developed a multiplier strategy to determine exhausted cell parameters for proliferation, secretion (TNFα/IFNγ), and CD8 T cell killing. Here we show an example for how these multipliers are utilized for the number of IRs for T cell proliferation (CD4 and CD8). CD4/CD8 T cell IFNγ secretion rate, CD4/CD8 T cell TNFα secretion rate, and CD8 T cell killing probability are calculated the same way. Cells with 3 IRs will have the most severe loss to their functional parameters. For example, if the probability of proliferation is sampled at 75% for the population of CD8 T cells then a single CD8 with 3 IRs will have a proliferation probability of no more than 75%*10% = 7.5%. For 2 IRs, it will be between 0% and 15%, and for 1 IR, 0% to 22.5%. We allow for heterogeneity by making each possible multiplier of IRs not just one multiplier but a range. This way not all cells with the same number of IRs are identical in their function. The ranges we selected also mean that a cell with a single IR could have a multiplier smaller than a cell with 3 IRs.

#### 2.2.8 HIV Infection

In our model, HIV is a diffusing entity where free virus spreads and degrades. Only activated CCR5+ CD4 T cells and macrophages can be infected probabilistically, either by free virus or cell-to-cell transmission (35,36). Immune cells randomly try free virus or cell-to-cell infection first, and if unsuccessful, the second method will be tried. For free virus infection, the virus concentration at the grid location of the cell needs to be at least 1 and a random number check against the probability of free virus infection needs to be successful. Cell-to-cell infection requires an infected neighbor and a successful probability of cell-to-cell infection check against a random number.

Infected cells enter an eclipse phase before they can productively shed virus; however, only activated cells can shed virus, at different rates for macrophages and CD4 T cells. If an activated HIV-infected CD4 T cell proliferates, then the progeny will also be HIV-infected(37). HIV-infected cells have an average lifespan of 2.2 days(38). We initialize the simulation with one large amount of virus (4.94e+04 ± 3.05e+04 [mean±std]) in the center of the grid. This ratio of virus to target cells (macrophages and CD4s) results in a large multiplicity of infection (MOI) (22 ± 22.6). While we recognize this relatively high MOI is not a common practice in *ex vivo* experimental HIV infections, we assume it is sufficient for our purposes here because experiments usually stimulate cells (with IL-2) before HIV infection due to the challenging nature of artificially creating HIV-infected cells(35,36). In the process of identifying parameters, we discovered that the low number of initially activated cells in our simulations was a major hurdle in having an infection establish itself at lower MOIs. However, to maintain the participant-specific nature of our simulations we do not inflate the number of activated cells and assumed a high MOI of virus to cells instead.

#### 2.2.9 Initial Conditions and First Estimate Parameter Ranges

**Table 1** in Supplement Materials shows a detailed list of the participant-specific markers incorporated into simulations. We exclude any participant from the full PIA study that was missing data for any of the immune markers in **Table 1** in Supplement Materials. Given the remaining participant-specific data (116 participants), we determine the absolute number of PBMCs per individual by subtracting the absolute neutrophil count from the white blood cell count (**WBC_x10.9.L** – **Absolute_Neu_x10.9.L**) to get the absolute number of PBMCs. Next, we divide the absolute number of lymphocytes (**Absolute_Lymph_x10.9.L**) and the absolute number of monocytes **(Absolute_Mono_x10.9.L**) by the absolute number of PBMCs we calculated to get the percentage of lymphocytes and monocytes per individual.

To assess the impact of IRs, we map model parameters to corresponding participant-specific markers where available (33 participant specific parameters) and sample other parameters within biologically feasible ranges (41 parameters). To find biologically feasible values for non-participant-specific parameters, we first used Latin Hypercube sampling (LHS)(39) on the parameters in **Table 3** marked with an asterisk. We generated 580 samples with 3 replicates each to give 5 unique parameter sets per participant in triplicate (1740 total runs). We ensure each participant has a baseline number of PBMCs that is high enough to cover all specific cell categories, except for triple exhausted CD4 or CD8 T cells. Using this LHS sampling approach with the strategy discussed in **2.2.7** and **Equation 1** for parameters affected by exhaustion, we selected biologically feasible parameter combinations by identifying simulations that reliably produced a robust range of HIV-infected macrophages and CD4 T cells and selected 20 representative parameter sets from these. We simulated these 20 parameter sets for each participant in triplicate for a total of 6960 runs (116 participants x 20 parameter sets x 3 replicates). **Table 3** shows the initial parameter ranges used to determine the 20 parameter sets in the analysis. The final 20 parameter sets are provided in Supplement Materials with the 41 parameters that are varied between sets and constant parameters. The full 116 participant parameters are provided in the Supplement as well. Analysis was completed on the average of the 3 replicates reducing the total simulation size to 2320. The size of each group is as follows: CHU_<5_=240, CHU_≥5_=400, CPHIV-VS_<5_=260, CPHIV-VS_≥5_=380, CPHIV- VA_<5_=480, and CPHIV-VA_≥5_=560. Two-tailed ttests were completed between each group for statistical analysis.

## 3. Results

### HIV status and age independently affect immune responses to new HIV exposure

To predict the overall impact of participant-specific immune profiles on infection progression, we simulate our virtual *in vitro* infections and quantify viral infection dynamics (**Fig. 3a,b**) and cellular death (**Fig. 3c**). We show Day 2-4 post-infection to evaluate HIV infection dynamics as this is the peak of HIV viral concentrations and infected cells. HIV-infected cell numbers and viral concentration rapidly decline after Day 4 and are nearly zero by Day 10.

**Fig. 3:**
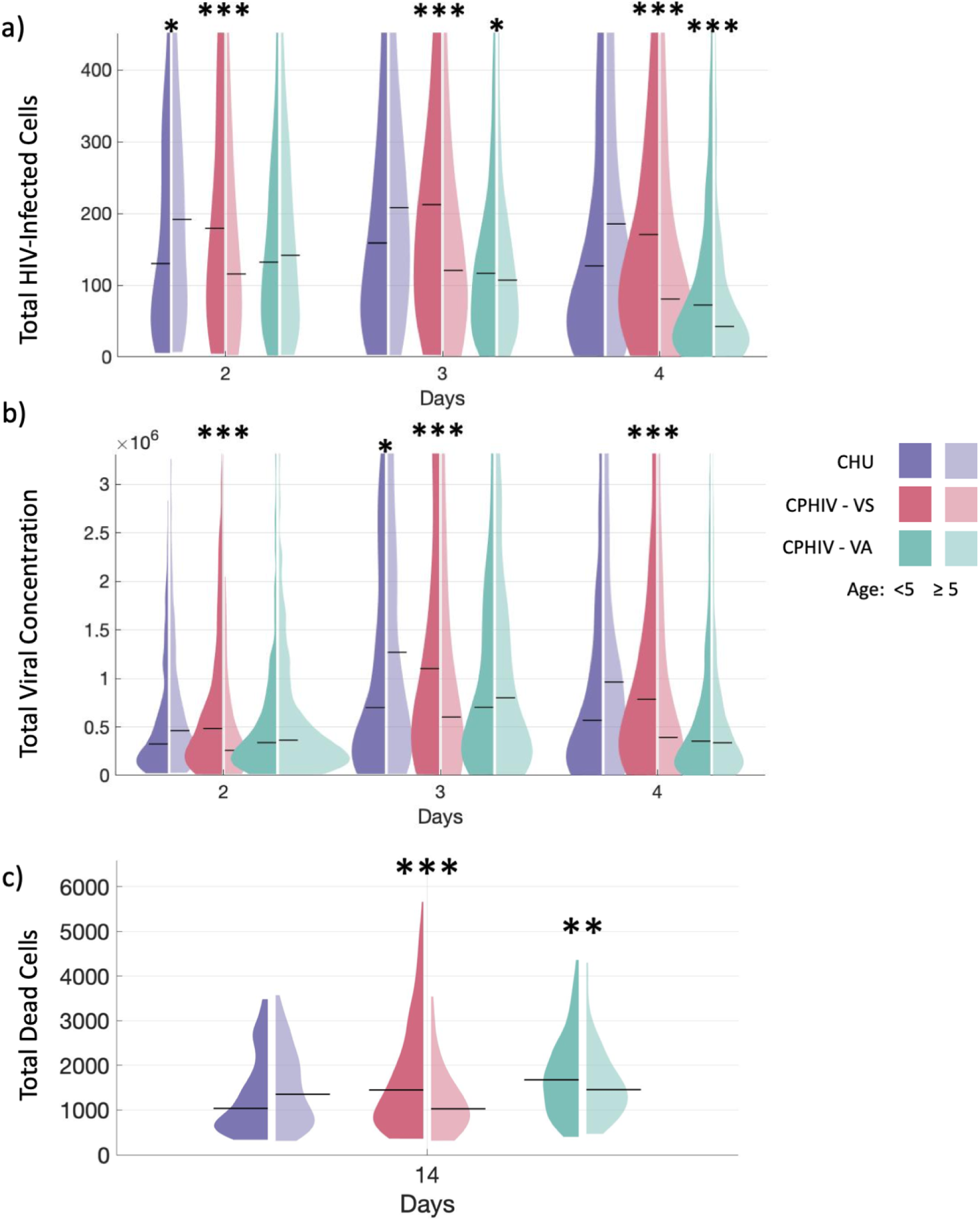
HIV status and age independently affect immune responses to new HIV exposure. a) Total HIV-infected cells for Days 2-4 post infection, b) total viral concentration for Days 2-4 post infection, c) total cell death at Day 14 post-infection. Left violin plots: participant age < 5, right: participant age ≥ 5. CHU: 0, CPHIV-VS: < 200 and CPHIV-VA: ≥ 200 HIV RNA copies/mL. From left to right n=240, 400, 260, 380, 480, 560. Violin plots show the distribution for each participant group, with black line indicating median. Y-axis for (a) and (b) are set to the 3^rd^ quartile for visibility, the full y range is shown in Fig S1 in Supplement Materials. ***p <= 1e-4, **p<=1e-3, *p<=0.01. Significance is determined with a two tailed t test between age groups for each viral participant group with α= 0.01 shown on the graph.

Cells from participants under 5 years of age (CHU_<5_) consistently have less virally infected cells, lower viral concentrations and less total cell death compared to cells from participants over 5 years of age (CHU_≥5_), however, majority are not significantly different (purple **Fig. 3a-c**). Conversely, CPHIV-VS shows statistically significant differences between age groups (pink **Fig. 3a-c**) (p < 1e-4, **Table 4**) in all of these metrics but with older participants (CPHIV-VS_≥5_) controlling new infection better. Cells from participants under 5 years of age (CPHIV-VS_<5_) are consistently predicted to have higher numbers of HIV-infected cells and viral concentration. Furthermore, this group exhibits a significantly higher number of cell death compared to CPHIV-VS_≥5_ together suggesting less effective immune responses to new viral exposure in younger compared to older CPHIV-VS participants. Patterns between age groups within CPHIV-VA vary across outputs and timepoints (green **Fig. 3a-c**) but the differences are small. These findings indicate that age-specific impacts vary between participant groups depending on HIV status and viral suppression. CPHIV-VS participants appear to have the largest differences in viral concentration and cell death between age groups.

**Table 4:**
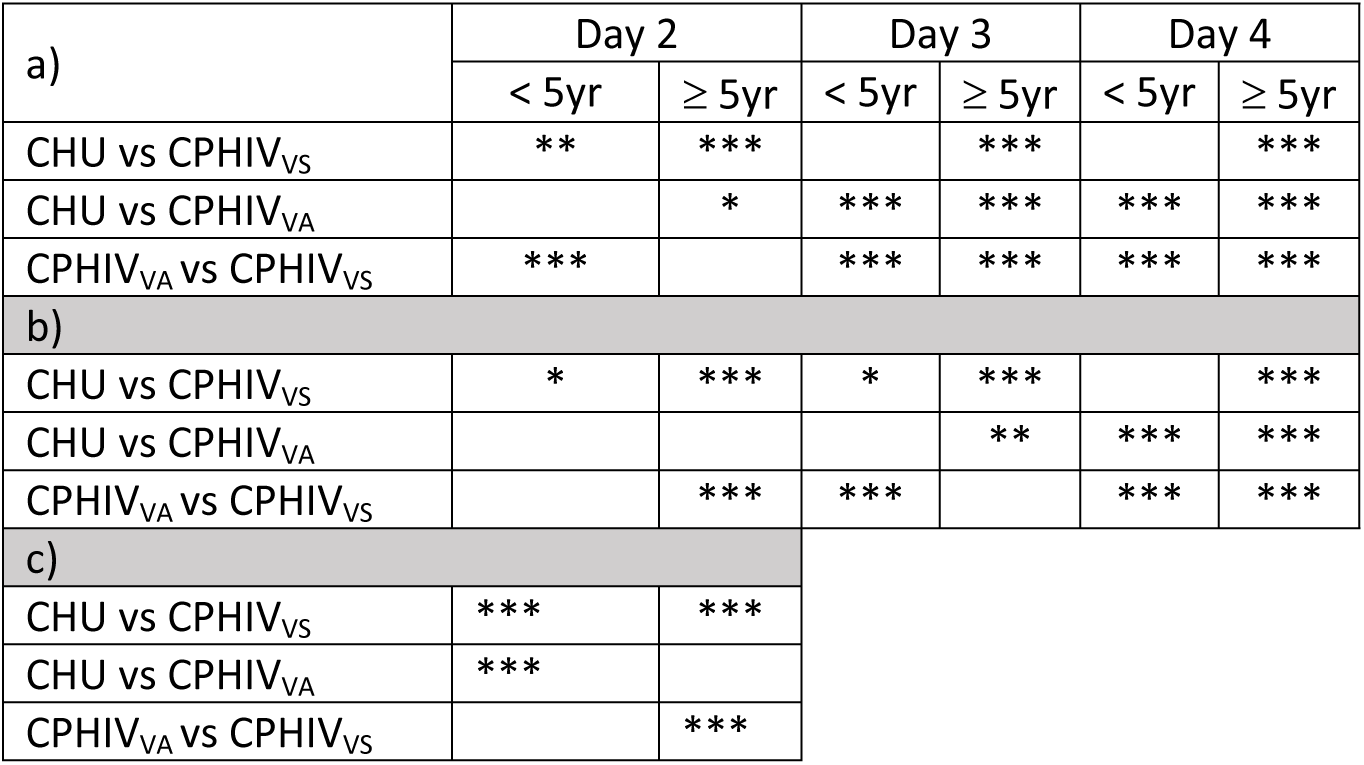
Statistical comparison between viral participant groups for each age participant group corresponding to Fig. 3a, b, and c respectively. Significance is determined with a two tailed t test between age-matched viral participant groups at α= 0.01. ***p <= 1e-4, **p<=1e-3, *p<=0.01.

For all participants under 5 years of age (darker left side of plots), the infection dynamics over a three-day period demonstrate a consistent trend (left plots **Fig. 3a-c**). CPHIV-VS_<5_ consistently has higher HIV-infected cells and virus concentration compared to CHU_<5_ and CPHIV-VA_<5_. CPHIV-VA_<5_ has lower viral load and HIV-infected cells (p < 1e-4, **Table 4**), and more cell death compared to CHU_<5_ (p < 1e-4, **Table 4**). This pattern suggests that for cells from participants under 5 years of age, CPHIV-VA_<5_ control viral loads to lower levels (**Fig. 3a,b**), but that this is associated with increased cell death (**Fig. 3c**). These patterns could indicate excessive inflammation which would impact the ability of CPHIV_<5_ to control infection longer term. In contrast, CPHIV-VS_<5_ simulations predict higher viral concentrations and infected cell numbers (**Fig. 3a,b**), but are also associated with higher cell death similar to CPHIV-VA_<5_ **Fig. 3c**. This suggests that there are key differences in the cell death mechanisms between virally suppressed and virally active CPHIV that result in different impacts on the viral concentration.

For participants over 5 years of age (lighter right side of plots), predictions show a more varied response **Fig. 3a-c**. Most groups are statistically significantly different from the other groups in number of infected cells, viral concentration and cell death (right violin plots in **Fig. 3 a-c**) (p < 1e-4, **Table 4**). For most time points, viral concentration and infected cells are lowest in CPHIV-VS_≥5_ and CPHIV-VA_≥5_ (**Fig. 3a,b)**, and cell death is lowest in CPHIV-VS_≥5_ (**Fig. 3c)**. These more varied patterns suggest more complex underlying mechanisms between the participant groups. However, our results indicate that cells from CPHIV-VS_≥5_ can maintain lower viral loads while also limiting cell death, which could reflect an effective short-term immune response.

### HIV status drives differential cell death patterns

While CPHIV-VS_<5_ and CPHIV-VA have increased predicted cell death compared to the other groups (**Fig. 3c**), these overall metrics could mask underlying differences in the types of cell death. A quantification of the cell types that contribute most to this overall cell death number reveals that CHU and CPHIV-VS have similar distributions: CD4 T cells make up the largest proportion of cell deaths (∼50%), followed by macrophages, then CD8 T cells (**Fig. 4a**). In contrast, in CPHIV-VA simulations, CD8 cells make up the largest proportion of cell deaths (**Fig. 4a**), almost 50% of all cell death compared to ∼25% or less for CHU and CPHIV-VS (p < 1-e4, **Table 2** in Supplement Materials). CHU_≥5_ and CPHIV-VA_≥5_ show reduced death of CD8 and CD4 T cells, respectively, compared to CHU_<5_ and CPHIV-VA_<5_ (**Fig. 4a**). This reduction in adaptive immune cell death is associated with an increase in macrophage death in both CHU_≥5_ and CPHIV- VA_≥5_ cohorts compared to their corresponding younger age groups. Thus, our simulations indicate that HIV status drives different cell death patterns by cell type, and that older age groups (CHU_≥5_ and CPHIV- VA_≥5_) have increased cell death in the innate immune cell populations.

**Fig. 4:**
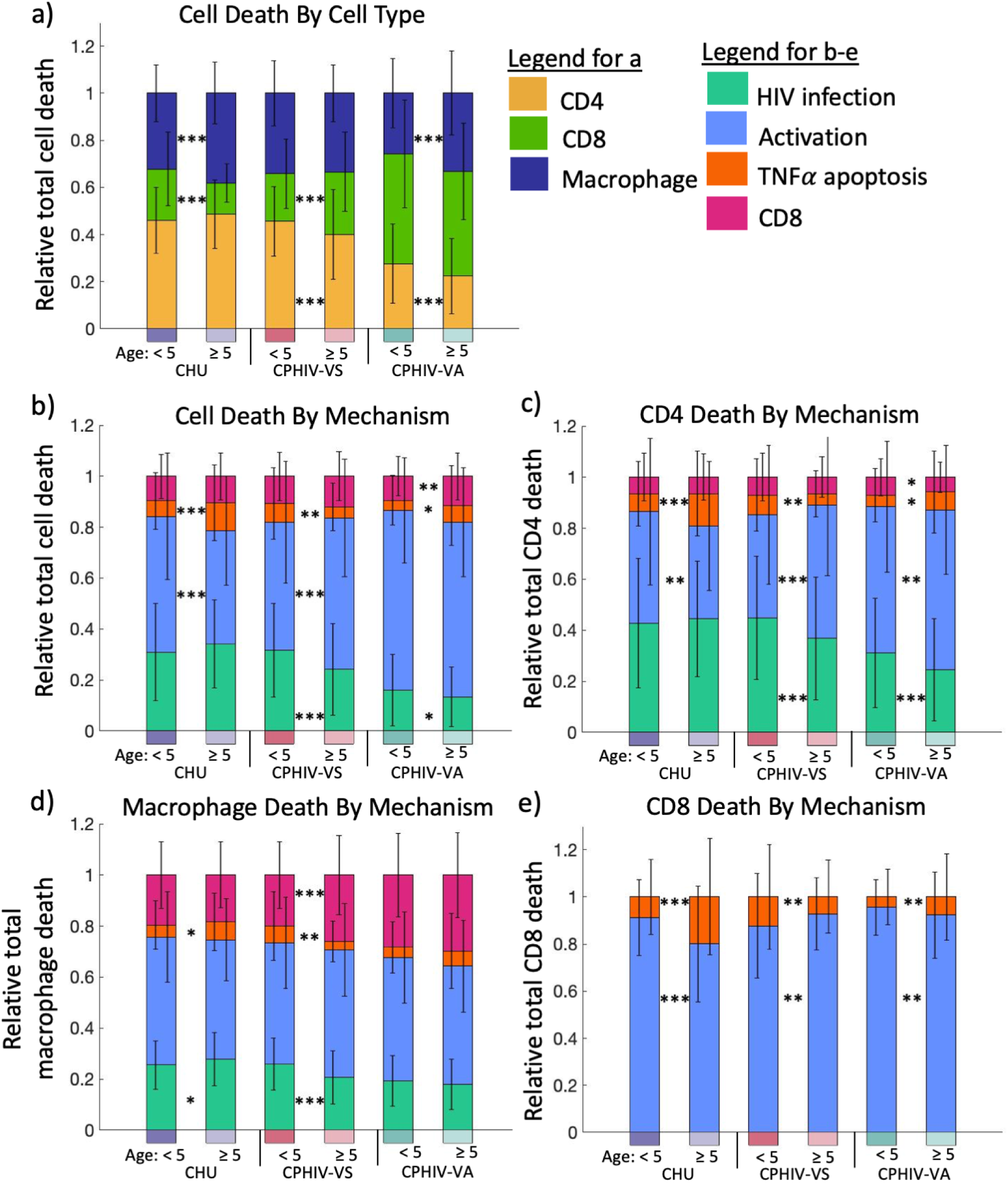
HIV status drives differential cell death patterns. a) Relative total death by cell type (bottom to top: CD4 T cell, macrophage, CD8 T cell). b) Relative total cell death by death mechanism (bottom to top: HIV infection, excessive activation, TNFα induced apoptosis and CD8 T cell killing). c) Relative CD4 T cell total death by mechanism (bottom to top: HIV infection, excessive activation, TNFα induced apoptosis and CD8 T cell killing). d) Relative macrophage cell death by mechanism (bottom to top: HIV infection, excessive activation, TNFα induced apoptosis and CD8 T cell killing). e) CD8 T cell death by mechanism (bottom to top: excessive activation, TNFα induced apoptosis). Timepoint = day 14. Bars display the mean for each group with error bars showing the standard deviation. CHU: 0, CPHIV-VS: < 200 and CPHIV-VA: ≥ 200 HIV RNA copies/mL. From left to right n=240, 400, 260, 380, 480, 560. ***p <= 1e-4, **p<=1e-3, *p<=0.01. Significance is determined with a two-tailed t test between age groups for each viral participant group with α= 0.01 shown on the graph and for age-matched groups given in Table 2 in Supplement Materials.

The proportions of cell death by different mechanisms (HIV infection, excessive activation, TNF-induced apoptosis or CD8 cytotoxic killing) provide further insight into the impacts of HIV status and age (**Fig. 4**b). Cell death from excessive activation (represented by Ki67+ and CD38+) is the dominant cell death mechanism in all participants groups. CHU and CPHIV-VS have significantly more deaths from HIV infection compared to CPHIV-VA (p < 1-e4, **Table 2** in Supplement Materials). CPHIV-VS shows the largest age-dependent differences between cell death mechanisms, with older participants having a larger proportion of cell deaths from excessive activation, and smaller proportions due to HIV infection and TNFα-induced apoptosis compared to the younger participants. This contrasts with CHU, where the older participants have a smaller proportion of cell deaths due to excessive activation compared to younger participants. Thus, while CPHIV-VS_<5_ have similar cell death numbers to CPHIV-VA (**Fig. 3c**), CPHIV-VS_≥5_ participants have similar cell death mechanism patterns to CPHIV-VA (**Fig. 4b**). These similarities in cell death mechanisms between CPHIV-VS_≥5_ and CPHIV-VA could help explain why these groups also have similar predicted viral load and infected cells (**Fig. 3a,b**), despite differences in cell death numbers (**Fig. 3c**).

An evaluation of cell death mechanisms for each immune cell type (**Fig. 4c**) further reinforces these observations and identifies how death mechanisms change for different cell types in each of the participant groups. Excessive activation remains the dominant cell death mechanism within each cell type, except for HIV-induced CD4 T cell death in CHU_≥5_ and CPHIV-VS_<5_. The overall pattern of high proportions of cell death from HIV infection in CHU and CPHIV-VS_<5_ (**Fig. 4b**) is present in both CD4 T cells and macrophages compared to CPHIV-VA (**Fig. 4c,d**) (p < 1-e4, **Table 2** in Supplement Materials). The age-dependent differences between CPHIV-VS are present in CD4 T cells, CD8 T cells and macrophages, with the most notable differences in excessive activation deaths in CD4 and CD8 T cells (again closely resembling patterns in CPHIV-VA).

Taken together, these results reveal unique patterns for each participant group both in which cell types are dying as well as how they are dying in response to short-term exposure to new HIV infection. These patterns are a result of the complex underlying composition of the immune cell population.

### Partially functional exhausted CD8 T cells contribute to anti-viral immune response

To better understand the role of exhausted CD8 T cells in the complex populations of immune cells from each of the participants, we quantify the contribution of exhausted CD8 T cells to key CD8 T cell functions during infection. Exhaustion affects CD8 T cell functions such as cytotoxic killing, proliferation, activation and TNFα and IFNγ secretion. Thus, we determined what proportion of each of these functions are performed by recently differentiated effector T cells, non-exhausted memory, and exhausted memory CD8 T cells in our simulations (**Fig. 5a-c**).

**Fig. 5:**
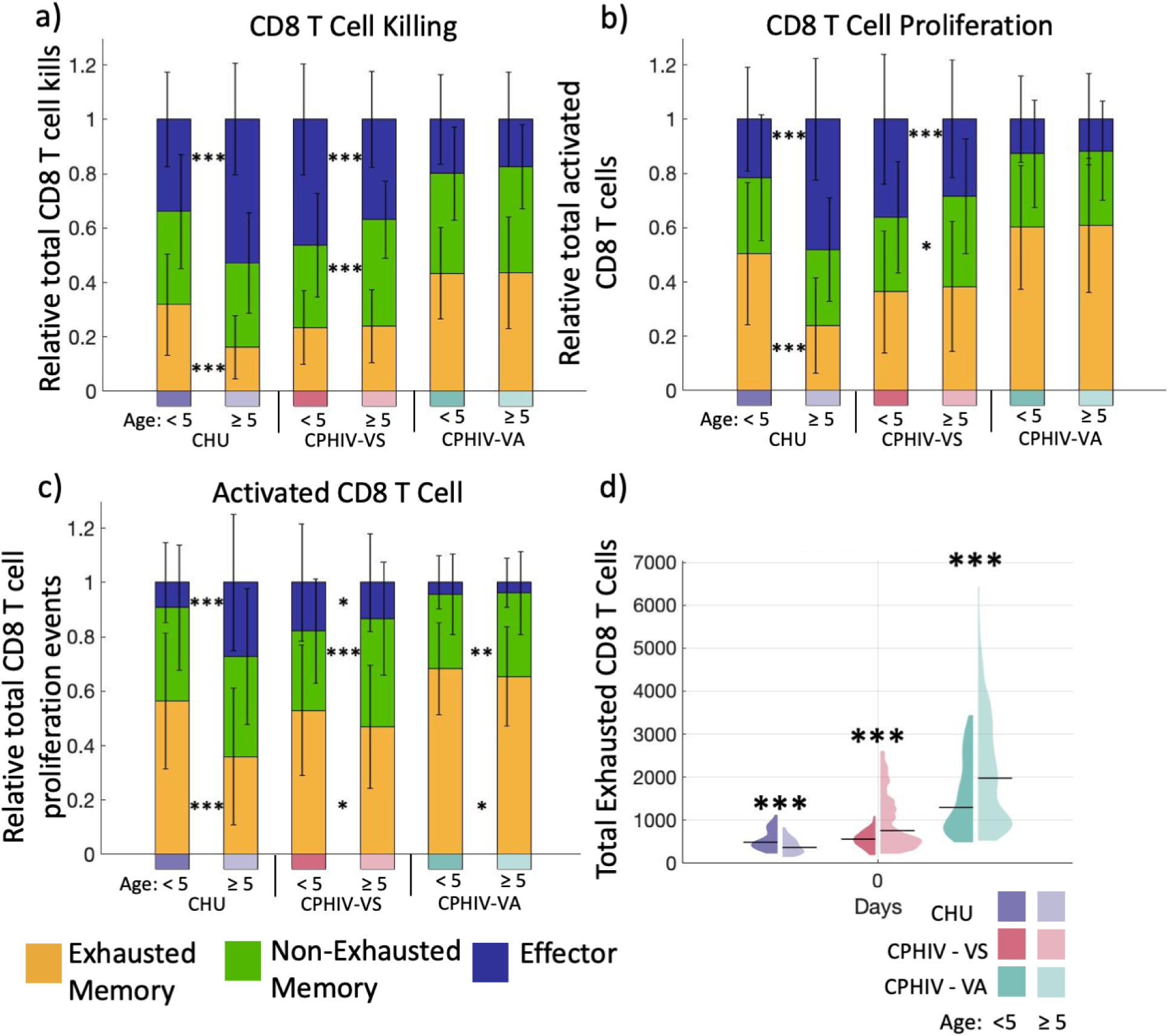
Partially functional exhausted CD8 T cells contribute to anti-viral immune response. a) Relative contributions by exhausted memory, non-exhausted memory, and recently differentiated effector CD8 T cells towards total CD8 T cell killing HIV-infected cells Day 14. b) Relative contributions by exhausted memory, non-exhausted memory, and recently differentiated effector CD8 T cells towards total CD8 T cell proliferation events by Day 14. c) Relative contributions by exhausted memory, non-exhausted memory, and recently differentiated effector CD8 T cells of total activated CD8 cells at Day 4. d) Total exhausted memory CD8 T cells at Day 0. Bars display the mean for each group with error bars showing the standard deviation. CHU: 0, CPHIV-VS: < 200 and CPHIV-VA: ≥ 200 HIV RNA copies/mL. From left to right n=240, 400, 260, 380, 480, 560. ***p <= 1e-4, **p<=1e-3, *p<=0.01. Significance is determined with a two-tailed t test between age groups for each viral participant group with α= 0.01 shown on the graph and for age-matched groups given in Table 3 in Supplement Materials. Violin plots show the distribution for each group and category with black line indicating median.

For CHU_≥5_ and CPHIV-VS_<5_, recently differentiated effector HIV-specific CD8 T cells kill the most HIV-infected cells (**Fig. 5a**). However, recently differentiated effector CD8 T cells contribute the least to CD8 T cell activation (**Fig. 5b**) for all age and viral concentration groups. In contrast, CPHIV-VA simulations show that the majority of CD8 T cell killing, proliferation and activation is from exhausted memory CD8 T cells (**Fig. 5a-c**). This indicates that while exhausted memory CD8 T cells are dominant in these CD8 T cell functions in CPHIV-VA simulations, the functions are more distributed among CD8 T cell subtypes in CHU and CPHIV-VS. However, it is important to note that the average number of total CD8 T cell proliferation events is less than 160 across all groups and the median is less than 25 across all groups with no significant differences between ages (**Fig. S2** in Supplement Materials). CHU_≥5_ consistently show that recently differentiated effector CD8 T cells contribute more (and exhausted cells contribute less) to killing (**Fig. 5a**), proliferation (**Fig. 5b**) and activation (**Fig. 5c**) as compared to the younger age group.

We believe that this counterintuitive prediction that the functionally impaired exhausted memory CD8 T cells are responsible for such a large proportion of CD8 T cell functions is related to the absolute numbers of exhausted cells. Indeed, CPHIV-VA simulations have higher numbers of exhausted CD8 T cells compared to CHU and CPHIV-VS (**Fig. 5d)** (p < 1-e4, **Table 3** in Supplement Materials), in agreement with the larger contribution of exhausted CD8 T cells to all CD8 T cell functions in CPHIV-VA (**Fig. 5a-c**) (p < 1- e4, **Table 3** in Supplement Materials). The opposite can be seen in CHU_≥5_ participants which have the fewest numbers of exhausted memory CD8 T cells compared to their younger participants at the start of the simulation (**Fig. 5d**), which coincides with the larger contribution of recently differentiated effector cells towards killing, proliferation and activation (**Fig. 5a-c**).

Taken together, our results indicate that despite significant T cell exhaustion in CPHIV-VA, these partially functional exhausted cells can continue to contribute to the anti-viral immune response during short-term infection.

## 4. Discussion

Our study, using computational models of short-term *in vitro* exposure of participant-specific cells to HIV, reveals counterintuitive findings regarding the potential contribution of exhausted T cells to anti-HIV immune responses in CHU and CPHIV. Cells from CPHIV-VA, despite showing the highest levels of exhaustion, exhibit lower model-predicted viral concentrations and fewer infected cells upon new HIV exposure compared to other groups. This unexpected result is accompanied by higher cell death rates, suggesting that short-term viral control may be linked to excessive inflammation, which can be potentially harmful in the long term. In contrast, cells from CPHIV-VS_≥5_ are predicted to maintain lower viral concentrations while limiting cell death. This could indicate a more effective and sustainable short-term immune response in CPHIV-VS_≥5_. CHU_≥5_ and CPHIV-VS_<5_ have the highest predicted levels of viral concentration and infected cells and high levels of cell death. Our model predicts distinct patterns of cell death across participant groups. CD8 T cell death contributes the most to cell death in CPHIV-VA while contributing the least in both age groups of CHU and CPHIV-VS which are dominated by CD4 T cell death. Cell death by excessive activation is the main death pathway in all groups, but HIV infection of CD4 T cells is much stronger in CHU and CPHIV_<5_ compared to other groups. Lastly, our results suggest that despite the impaired functionality of exhausted CD8 T cells (in proliferation, cytokine secretion and killing), large enough numbers of exhausted CD8 T cells can overcome these limitations. Our simulations therefore highlight unique patterns of immune responses to short-term HIV exposure resulting from unique and complex participant-specific immune profiles. We identify immune response patterns that vary with both age and HIV status.

Simulations for CPHIV-VS_<5_ participants predict higher viral concentration and cell death as compared to older participants who are virally suppressed. This suggests that CPHIV-VS_<5y_ are mounting an ineffective inflammatory response to the new HIV exposure. This is consistent with evolving immunologic maturity over the first 5 years of life (60–62). Neonatal monocytes have reduced production of pro-inflammatory cytokines compared to adult monocytes, including TNFα and IFN-γ (63–66). TNF-α remains low in infants up until age 5 years(67) then increases until it reaches peak concentrations at age 14 years that are higher than adult levels, then drops to adult levels(68). IFN-γ was seen to be in normal adult range from age 3-17 years(68) which leads to the general assumption that increased levels indicate inflammation. Overall, CPHIV-VS_<5y_ immune response patterns strongly resemble the age-matched healthy control group of CHU. Our simulations suggest that CPHIV who become virally suppressed before they reach 5 years in age, have limited immune exhaustion which enables them to respond to new infection challenges in a similar manner to children without HIV.

In contrast, cells from CPHIV-VS_≥5y_ are predicted to have lower viral load and lower cell death compared to their younger counterparts as well as CPHIV-VA. This suggests that CPHIV-VS_≥5y_ are able to mount an effective immune response that can limit viral replication with less host cell death. However, in our analysis this immune response is associated with more cell death from excessive activation rather than cytotoxic killing of infected cells. Thus, it appears that cells from CPHIV-VS_≥5y_ can indeed mount a more inflammatory response (more immune cell activation), but that this is not necessarily associated with improved killing of infected cells. This observation may be one mechanism that contributes to chronic inflammation reported in longer-term infections. Viral suppression in infants and young children under 5 years initiating ART takes longer than in older adults, which has been attributed to immunologic immaturity (69,70). In a Canadian cohort, children above age 5 years starting ART had a higher probability of viral rebound, which has also been seen in other cohorts of children (71,72). Indeed, the cell death patterns for CPHIV-VS_≥5y_ resemble those from CPHIV-VA more closely than CPHIV-VS_<5_. This suggests that despite viral suppression, the length of time since infection in CPHIV-VS_≥5y_ participants has been long enough that the cell death mechanisms have started to shift more closely to virally active patterns, even though the metrics of exhaustion still show much lower levels compared to virally active participants.

Finally, CPHIV-VA show very little differences between age groups, suggesting that the impacts of active viral replication are larger than the impacts of age. Our results indicate that in the short-term, viral replication is limited due to large amounts of cell death that mostly stems from excessive activation of macrophages, CD4 and CD8 T cells. Many forms of programmed cell death have been associated with HIV infection, including apoptotic and nonapoptotic pathways(73). The induction of apoptosis with pro-apoptotic TNF-derived peptides is associated with reduced HIV-associated cell death and reduced viral load(74). Meanwhile, nonapoptotic pathways, like necroptosis and pyroptosis, are thought to be host-detrimental leading to increased cell death and immunosuppression(73). Activation-induced cell death of T cells is found to be higher in persons with HIV, compared to healthy controls, with a reduction following ART (75–78). This evidence supports simulation results that CPHIV have more activation-associated deaths compared to CHU, and CPHIV-VS has fewer activation-induced deaths than CPHIV-VA. Surprisingly, our simulations predict that a sizable portion of exhausted CD8 T cells continue to contribute to CD8 T cell cytotoxic killing, proliferation, and cytokine secretion in these short-term infections. However, the large amounts of cell death from excessive activation indicate that these exhausted cells are likely to be depleted as the infection progresses to longer periods, possibly leading to increased viral replication long-term.

As with all computational models, there are limitations associated with necessary simplifying assumptions. We acknowledge that our short-term simulations of exposure to new HIV infections (emulating *in vitro* infection assays) are limited to making predictions about short-term *in vitro* dynamics, and therefore do not account for other dynamics that emerge *in vivo* over longer periods. Future work will incorporate methods presented here into simulations of longer-term *in vitro* experiments including periodic cell supplementations, and into our existing models of *in vivo* infections(79,80). This will inform both *in vitro* experimental design as well as understanding of *in vivo* infection dynamics. Additional limitations include necessary assumptions about how to map the IR markers to simulation mechanisms, including that IR expression is linearly correlated with function impairment as well as having any ability at all to perform a function and which functions are impacted by IR expression. We believe that these assumptions are reasonable given current knowledge (5,33), and can be adapted as more data on the impact of individual IR markers become available.

In short, our computational simulations integrating individualized immune profiles with known mechanisms and parameters allows us to predict the combined outcome of complex immune responses to new HIV exposure. Our simulations suggest that partially-functional exhausted CD8+ T cells can contribute to short-term viral suppression in CPHIV. Further, our results indicate that viral suppression in CPHIV can allow immune responses similar to CHU, but only in CPHIV_<5_. These findings could inform immunotherapeutic strategies, HIV cure studies, and HIV early treatment studies.

## Ethics Statement

Ethical approval for this study was obtained from New York University (10-02586) and Kenyatta National Hospital/ University of Nairobi (P283/07/2011). The studies were conducted in accordance with the local legislation and institutional requirements. Written informed consent for participation in this study was provided by the participants’ legal guardians/next of kin and verbal assent from participants older than 7 years.

## Data Availability

Supplemental material, including participant-specific and non-participant specific parameters, MATLAB data and graphing scripts, are archived on Zenodo (10.5281/zenodo.17148547). The Repast model and accompanying parameter file to run the code can be found at: https://github.itap.purdue.edu/ElsjePienaarGroup/PediatricHIVExhaustionModel.

